# Psilocybin decreases reward-seeking behavior accompanied by increased activity of parvalbumin neurons with perineuronal nets in the medial prefrontal cortex

**DOI:** 10.64898/2025.12.22.696123

**Authors:** Jenna Houff, Andrew Williams, Obie Allen, Barbara Gisabella, Harry Pantazopoulos, Alberto Del Arco

## Abstract

Clinical trials suggest that a single dose of psilocybin is an effective treatment for substance use disorders (SUDs). Choice impulsivity is a value-based decision-making bias that predicts drug-intake escalation and is commonly associated with SUDs. The dorsomedial prefrontal cortex (dmPFC) regulates choice impulsivity and is enriched with 5-HT_2A_ receptors that mediate effects of psilocybin. We hypothesized that psilocybin has long-term (≥48 hours) effects on choice impulsivity in association with dmPFC inhibitory interneurons with perineuronal nets (PNNs). Male Long Evans rats were trained in a delay discounting task (DDT) where rats chose between delayed large rewards (LR) and immediate small rewards (SR). 48 hours after psilocybin or vehicle injections, DDT was assessed, and rats’ brains processed for microscopy analysis of extracellular matrix (PNNs) together with inhibitory parvalbumin (PV) interneurons and c-fos as a marker of neuronal activity. Psilocybin acutely increased head-twitch responses. Psilocybin decreased LR choices and increased the latency to LR choices 48 hours after administration. These effects were independent of delay and therefore not consistent with changes in impulsivity. Psilocybin also increased the density of PNN+PV+cFos triple-labeled neurons in the dmPFC. These results suggest that psilocybin decreases reward seeking through the increased activation of dmPFC PV interneurons with PNNs.

## INTRODUCTION

Recent clinical and preclinical studies suggest that the psychedelic compound psilocybin is a promising treatment for substance use disorders (SUDs) (DuPont and Johnson 2025; Li et al. 2025). For example, psilocybin administered together with psychological support (i.e., psilocybin-assisted psychotherapy) decreases the consumption of nicotine and alcohol in humans (Johnson et al. 2017; Vargas et al. 2021; Bogenschutz et al. 2022) and decreases the consumption of alcohol (Jeanblanc et al. 2024) as well as the risk for relapse after heroin and alcohol self-administration in animal models (Meinhardt et al. 2021; Floris et al. 2025). Importantly, these studies emphasize that one single dose of psilocybin is sufficient to produce long-lasting therapeutic and/or behavioral effects (Goodwin et al. 2022; Knudsen 2023). However, the mechanisms by which a single dose of psilocybin results in long-term modification of brain processes that regulate reward-seeking behavior are still unknown.

Impulsivity is strongly associated with substance use and the risk of developing SUDs (Volkow and Baler 2015; Kozak et al. 2019). Choice impulsivity is a value-based decision-making bias that leads individuals (i.e., humans and experimental animals) to overvalue immediate small reward choices compared to large rewards that come with a time delay (Frost and McNaughton 2017; Levitt et al. 2020). By using the delay-discounting task (DDT), previous studies have shown that increased impulsivity (i.e., decreased preference for delayed rewards) anticipates the escalation of drug use and relapse, and is a common symptom in psychiatric disorders including SUDs (Dalley et al. 2011; Hamilton et al. 2015; Volkow and Baler 2015; Amlung et al. 2019). The dorsomedial prefrontal cortex (dmPFC) encodes reward cues (Moorman and Aston-Jones 2015; Otis et al. 2017; Harris et al. 2025) and regulates value-based decision-making behavior (Bercovici et al. 2023; Wenzel et al. 2023). Notably, the dmPFC is rich in 5-HT_2A_ serotonin receptors which are a key cellular target mediating the actions of classic psychedelic drugs such as psilocybin (Cameron et al. 2023; Vargas et al. 2023; Shao et al. 2025). Specifically, both pyramidal neurons and fast-firing inhibitory interneurons (i.e., parvalbumin [PV] interneurons) in deep layers of the prefrontal cortex have a high expression of these receptors (Willins and Meltzer 1997; De Almeida and Mengod 2007; Andrade 2011). Here we hypothesized that psilocybin decreases choice impulsivity through remodeling the brain extracellular matrix and PV interneurons activity in the dmPFC.

By administering 5-HT_2A_ agonists and antagonists in rats, previous studies suggest that the activation of 5-HT_2A_ receptors acutely increases impulsivity assessed through the one- or five-choice reaction time task (Sholler et al. 2019; Higgins et al. 2021). In contrast, by using the DDT, a recent study reported no changes in choice impulsivity after psilocybin injections (Roberts et al. 2023) and, therefore, whether psilocybin modulates impulsivity remains unclear. Importantly, the above-mentioned studies focused on the acute effects (i.e., ≤ 24 hours) of psilocybin or 5-HT_2A_ compounds and therefore do not provide information about potential long-term (i.e., ≥ 48 hours) effects of psilocybin on processing reward cues. To determine whether psilocybin produces long-term effects on reward processing is highly relevant in the context of its potential use as a treatment for SUDs. Recent studies reported key plastic changes in the dmPFC, such as increases in the number and size of dendritic spines, that peak at 24-48 hours and last several days after psilocybin administration (Shao et al. 2021, 2025). Based on this evidence, in the current study we tested the effects of psilocybin on DDT performance as well as perineuronal nets (PNNs) and PV interneurons in the dmPFC 48 hours after administration in order to examine long-term rather than the initial effects of this psychedelic drug.

PNNs are specialized extracellular matrix structures that regulate experience-dependent neural plasticity and neuronal firing. Alterations of PNNs are linked to long-term behavioral changes associated with substance use in both animal models and humans (Slaker et al. 2018; Brown and Sorg 2022; Valeri et al. 2023). Recent studies have found that psilocybin and other psychedelic drugs increase the expression of genes involved in PNN remodeling and suggest that PNNs are linked to the therapeutic benefits produced by these drugs (Ly et al. 2018; Nardou et al. 2023; Huang et al. 2024). Notably, it has been shown that PV inhibitory interneurons in the dmPFC, which PNNs typically surround, are enriched with 5-HT_2A_ serotonin receptors (Willins and Meltzer 1997; De Almeida and Mengod 2007; Andrade 2011; Hsiao et al. 2025) and that the effects of psilocybin on PNNs depends on these receptors (Ly et al. 2018; Huang et al. 2024). PNNs regulate the activity of PV interneurons in the dmPFC (Slaker et al. 2018; Fawcett et al. 2019), which in turn can facilitate the extinction of reward-seeking behavior (Sparta et al. 2014).

The aim of this study is to determine whether a single injection of psilocybin modulates value-based reward-seeking behavior and whether this modulation is associated with changes in PNNs and activity of PV interneurons in the dmPFC. To this end, we assess performance in the DDT, the density of PNNs and the activity of PV interneurons through cFos labelling 48 hours after psilocybin (1 mg/kg, i.p.) administration in order to examine the sustained effects rather than the immediate effects of psilocybin. This dose of psilocybin reliably produces behavioral and plastic effects, including the expression of Fos in the dmPFC of rats (Shao et al. 2021, 2025; Davoudian et al. 2023; Funk et al. 2024; Anderson and Robinson 2025). We also evaluate these markers in the ventromedial prefrontal cortex (vmPFC) since some studies suggest that dmPFC and vmPFC play different roles in reward-seeking behavior (Peters et al. 2009; Burgos-Robles et al. 2013; Moorman et al. 2015).

## METHODS

### Animals

18 adult male Long Evans rats (Envigo, Indianapolis, IN) (350–375 g) were pair-housed in standard polycarbonate cages (45 x 24 x 20 cm) on a 12 hours light/dark cycle (lights on at 9:00 P.M.). The sample size was based on our previous work (Martinez et al. 2024). All experiments were performed during the dark phase when the animals are most active. The rats were placed on a mild food-restricted diet (15 g of chow per rat and day) two days before starting behavioral experiments. All procedures were approved by the University of Mississippi Institutional Animal Review Board and were conducted in accordance with the National Institute of Health Guide for the Care and Use of Laboratory Animals.

### Experimental design

Two weeks after their arrival at the animal facility, all rats were handled, for at least three days, and then habituated to the operant chambers and trained in the DDT (Figure 1). The experimental timeline is represented in Figure 2a. Once animals achieved stable performance, they were divided into two groups: Vehicle (n= 8) and Psilocybin (n= 8); two animals that did not perform the task were discarded. After training in the DDT, rats were taken to a different room; the Psilocybin group was administered a single dose of psilocybin (1 mg/kg, i.p.) and the Vehicle group 1 ml/kg of vehicle (saline). Head-twitch responses for 60 min were assessed immediately after injections. Then, 24 and 48 hours after psilocybin or vehicle injections, both groups of rats were tested in the DDT. Finally, 75-90 min after the last DDT session, the animals were euthanized, and their brains extracted for immunohistochemistry assays.

**Figure 1:**
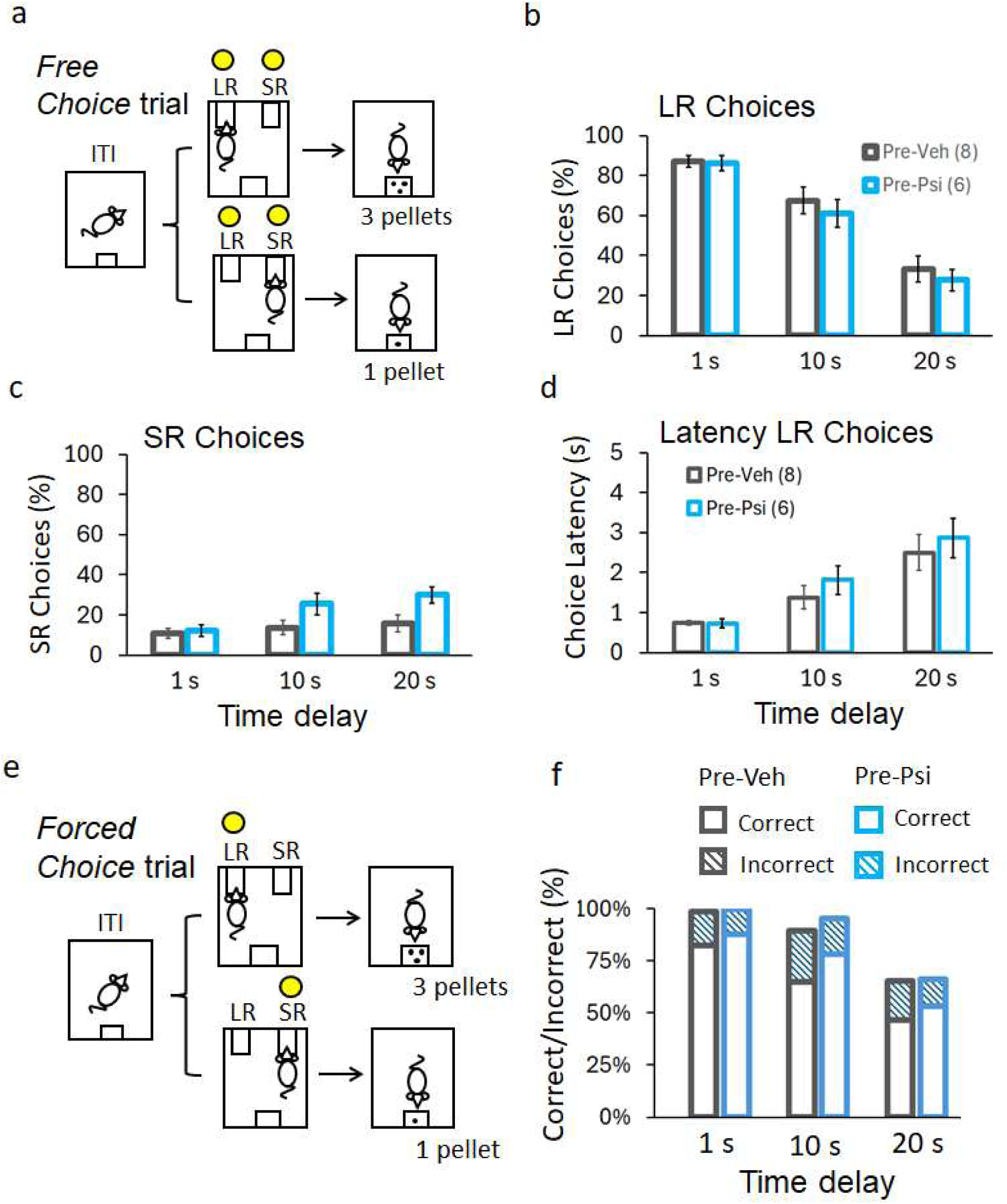
Delay discounting task and behavior before psilocybin (or vehicle) injections. In *Free Choice* trials **(a)**, the two levers are active, and rats need to make a choice. LR choices (**b**), SR choices (**c**) and choice latency (**d**) are evaluated at different delays for LR (1 s, 10 s and 20 s) during Free Choice trials. In *Forced Choice* trials **(e)**, the two levers, LR and SR, are extended but only the active lever shows a cue light. Accuracy (correct and incorrect responses) is evaluated at different delays for LR (1 s, 10 s and 20 s) during Forced Choice trials **(f)**. The delay for SR was always 1 s. Data are the mean ±SEM. The mean values are the average for the last three training sessions for every dependent variable at different delays.

**Figure 2:**
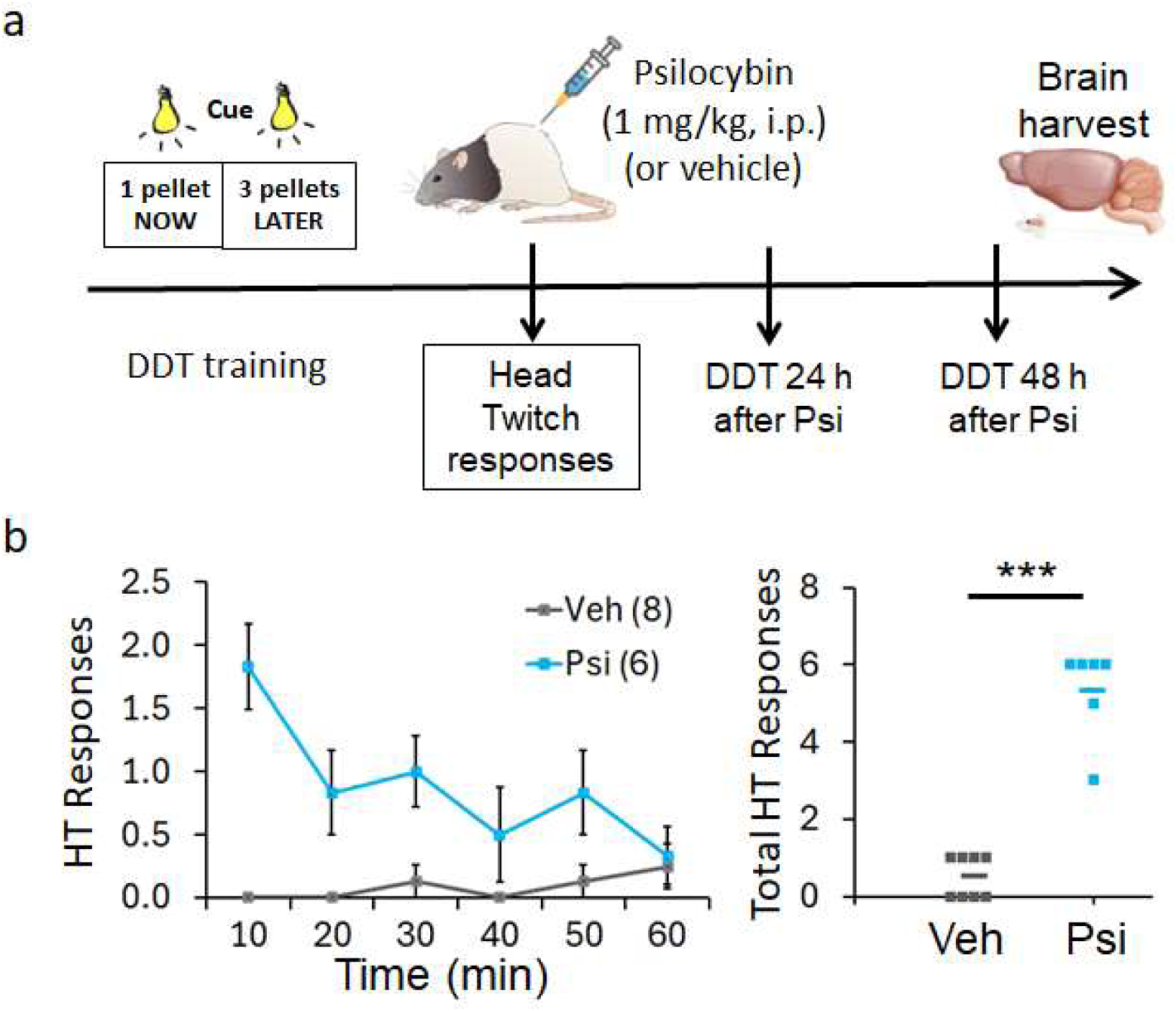
Timeline for the experimental protocol and head twitch (HT) responses **(a)**. Rats were trained in the delay discounting task (DDT) until stable performance. Rats received a single injection of psilocybin (or vehicle) and HT responses were immediately assessed. 24 hours and 48 hours after injections, rats were tested in the DDT. 75-90 minutes after the last DDT session, rats were deeply anesthetized and the brains extracted for immunohistochemical analysis. **(b)** Graphs show the temporal profile of HT responses and the total number of HT responses for 60 min in both groups, Vehicle and Psilocybin. Data are mean ±SEM. Dots represent individual animals. *** p< 0.001 compared to Vehicle after independent *t* test.

### Delay-discounting task (DDT)

We utilized the DDT used in our previous studies (Martinez et al. 2024). The apparatus consisted of a sound attenuated operant chamber with two retractable levers (right and left) and a food trough in between them in the same wall (Coulbourn instruments). The chamber contained a house light that was on during the entire session. First, animals were trained to establish the lever–response contingency through a reward-shaping procedure using both levers (3-4 days) (dustless sugar pellets, 45mg; Bio-Serv). Second, they were trained to discriminate between a large-reward (LR) lever and a small-reward (SR) lever (FR1, 30 min sessions). Pressing the LR lever delivered 3 sugar pellets while pressing the SR delivered 1 sugar pellet. This training phase lasted until they pressed the SR lever <25% of the trials (5-6 days). Then, the DDT training started. The task consisted of 3 blocks of 20 trials each (60 trials total, < 60 min sessions). In each block, the first 10 trials were *Force Choice* trials (Figure 1e) in which the two levers were extended but the associated cue was lit in only one of them (one active lever). The right lever was associated with a SR (1 sugar pellet) and the left lever was associated with a LR (3 sugar pellets). This designation was counterbalanced among animals. The remaining 10 trials were *Free Choice* trials (Figure 1a) in which both levers were extended with the cue lights (two active levers), and animals were required to make a choice. Both levers were extended to a maximum of 10 s and were retracted after animals made their choice. The three blocks were different in the delay to receive the large reward after pressing the LR lever: 1 s (block 1), 10 s (block 2) and 20 s (block 3). The pellet/s were delivered to the food trough, which light was turned on then. If animals did not press any lever within the 10 s both levers were retracted, and the trial was considered an omission. Animals were trained for 12 days until stable performance, which typically was accomplished after 5-7 days of training. A two-way ANOVA with repeated measures with group (Pre-Vehicle, Pre-Psilocybin) as between subjects, and delay (1 s, 10 s and 20 s) and sessions (6 last training sessions) as within subjects was performed to confirm stable performance.

### Psilocybin injections and head twitch responses

One or two days after stable performance in the DDT, rats received a single injection of vehicle (saline, 1 ml/kg, i.p.) or psilocybin (1 mg/kg, i.p., Sigma-Aldrich). To assess head-twitch responses, rats were taken to a procedural room and acclimated for 30 min. Then, rats were injected with vehicle or psilocybin and then immediately placed in acrylic boxes (16” W x 16” D x 15” H) and video recorded for 60 min using a camera positioned above the box. Two trained observers blind to treatment live counted the number of head twitches per 5 min time stamps. The final co-occurrence between observers was above 90% and confirmed by video recordings. After the assessment, rats were brought back to their home cages. All injections were carried out between 10 am and 12 pm.

### Immunohistochemistry

75-90 min after the last DDT session, the animals were deeply anesthetized (isoflurane 5%) and then perfused with phosphate buffer (0.1 M) and 10% formalin. The brains were extracted, further fixed for 24 hours, and then placed in sucrose 30% and stored in the fridge until slicing.

Serial free-floating tissue sections (spanning level 5 to level 10 of the Swanson Brain Maps 3.0 rat brain atlas (http://larrywswanson.com/?page_id=164) were carried through antigen retrieval in citric acid buffer (0.1 M citric acid, 0.2 M Na2HPO4) heated to 80 °C for 30 minutes and incubated in primary antibody monoclonal mouse anti-PV (1:4,000; P3088; clone PARV-19, ascites fluids; Sigma-Aldrich, St. Louis, Missouri), biotinylated Wisteria floribunda agglutinin (WFA) lectin (1:1000, catalog #B-1355, Vector Labs), and rabbit anti-cFos (1:300; 9F6, cat#2250S) for 48 hours at 4 °C. Immunoblot characterization for anti-PV P3088 showed a single band corresponding to 12 kD (information kindly provided by Sigma-Aldrich). WFA is an established marker for PNNs that binds to non-sulfated N-acetyl-D-galactosamine residues on the terminal ends of CSPGs.

Sections were subsequently in secondary antibodies Alexa Fluor donkey anti-mouse 647 (1:300; cat# A31571; Alexa Fluor streptavidin 488; cat# S11223; and Alexa Fluor goat anti-rabbit 555; cat#A32732; Invitrogen, Carlsbard, CA) for 3 hours. All solutions were made in PBS with 0.2% Triton X (PBS-Tx) unless otherwise specified. Immunostained sections were mounted on gelatin-coated glass slides, coverslipped with Dako Fluoromount fluorescence mounting medium (cat#S3023, Agilent Technologies) and coded for blinded quantitative analysis. All sections included in the study were processed simultaneously within the same session to avoid procedural differences.

Quantification was performed using an Olympus BX61 fluorescent microscope interfaced with Stereo-Investigator v11. The superficial and deep layers of the medial prefrontal cortex were traced for an area measurement using a 10X objective, and PV neurons, WFA-labeled PNNs and cFos positive neurons were quantified on 40X magnification using Stereo-Investigator v11.

### Data analysis

Two- and three-way ANOVAs with repeated measures with group (Vehicle, Psilocybin) as between subjects, and delay (1, 10 and 20 s) and/or session (24 hours and 48 hours) as within subjects were performed to compare the following dependent variables: LR choice, SR choice, accuracy (i.e., number of correct and incorrect responses in forced choice trials), latency and omissions. Pair-wise planned comparisons were performed to assess group effects. Numerical densities of immunoreactive cells were calculated as Nd= ∑N / ∑V where N = sum of all cells counted in each region, and V is the volume of each region, calculated as V= ∑a • z, where *z* is the thickness of each section (30 µm) and *a* is area in µm^2^. Densities were transformed to logarithms prior to data statistical comparison between Vehicle and Psilocybin groups. A Student independent *t* test was used to evaluate the effects of psilocybin on head twitch responses and numerical densities of PNNs, PVs and cFos labelling in the medial prefrontal cortex. Pearson’s linear regression analysis was performed to assess correlation between triple labelling and LR choices (average of all delays) 48 hours after vehicle and psilocybin injections. Statistical analysis was performed with SPSS software, and the statistical significance was set at p< 0.05.

## RESULTS

### Delayed Discounting Behavior

Animals were trained in the DDT (**Figure 1a, e**) and then split into two groups, Pre-Vehicle and Pre-Psilocybin. Stable performance was confirmed by two-way ANOVA analysis (see Methods) that showed no differences in large reward (LR) choices among the last six training sessions (session: F_(5,60)_= 0.84, p= 0.493, η_p_^2^= 0.06) considering group (session x group: F_(5,60)_= 0.21, p= 0.957, η_p_^2^= 0.01) and delay (delay x session: F_(10,120)_= 0.70, p= 0.720, η_p_^2^= 0.05) as factors. As in our previous studies (Martinez et al. 2024), the number of LR and small reward (SR) choices (i.e., percentage of LR and SR lever presses) and the latency to press the LR lever (i.e., choice latency), were evaluated during the Free Choice trials. The number of correct and incorrect responses (i.e., accuracy) and the latency to press the lever were evaluated during Force Choice trials.

Figure 1 shows that all rats decrease their preference for LR and increase their preference for SR choices as the delay to LR delivery increases to 10 s and 20 s. Considering the average of the three last training sessions, a two-way ANOVA with repeated measures with group (Pre-Vehicle, Pre-Psilocybin) as the between subjects variable and delay (1 s, 10 s and 20 s) as the within subjects variable was used to analyze performance in the DDT. The results of this analysis revealed a decrease in the number of LR choices (**Figure 1b**) (delay: F_(2,24)_= 74.13, p< 0.001, η_p_^2^= 0.86), and an increase in the number of SR choices (**Figure 2c**) (delay: F_(2,24)_= 14.32, p< 0.001, η_p_^2^= 0.54) with higher delays. We also observed an increase in choice latency (**Figure 1d**) (delay: F_(2,24)_= 30.04, p< 0.001, η_p_^2^= 0.71) and a decrease in the number of correct responses in forced choice trials (**Figure 1f**) (delay: F_(2,24)_= 43.19, p< 0.001, η_p_^2^= 0.78) with higher delays. The number of incorrect responses in forced choice trials was not altered with higher delays (delay: F_(2,24)_= 2.64, p= 0.092, η_p_^2^= 0.18). No significant differences were found between Pre-Vehicle and Pre-Psilocybin groups in any of these behavioral parameters.

### Effects of psilocybin on head-twitch responses

After stable performance in the DDT, rats were injected once with vehicle (saline, 1 ml/kg, i.p.) or psilocybin (1 mg/kg, i.p.) (**Figure 2a**). Due to complications during the injection, two rats were not injected with psilocybin, and one rat received 0.33 mg/kg instead of 1 mg/kg of psilocybin. This rat, however, was included in the Psilocybin group based on performance [head twitch= 6 (Group min= 3 and max= 6); density triple labelling= 2.78 (Group min=2.16 and max= 7.99); LR choices at 48 hours= 20 (Group min= 0 and max= 60)]. We assessed head-twitch responses immediately after injections. As shown in **Figure 2b**, psilocybin increased the total number of head-twitch responses in 60 min compared to vehicle (Vehicle= 0.5 ±0.2; Psilocybin= 5.33 ±0.54; t(12)= 10.14. p< 0.001). This effect was maximal 10 min after administration and returned to baseline after 60 min.

### Effects of psilocybin on delay discounting behavior

Rats were tested in the DDT 24 and 48 hours after psilocybin injections. To analyze whether psilocybin alters delay discounting behavior, three-way ANOVAs with repeated measures were performed, considering group (Vehicle, Psilocybin) as the between subjects’ variable and delay (1 s, 10 s and 20 s) and sessions (24 and 48 hours) as within subjects’ variables, to compare LR choices, SR choices, accuracy, latencies and omissions. **Figure 3**, **Figure 4** and supplemental **Figure 1** display the effects of psilocybin on these behavioral parameters 24 and 48 hours after injections compared to baseline (before psilocybin injections).

**Figure 3:**
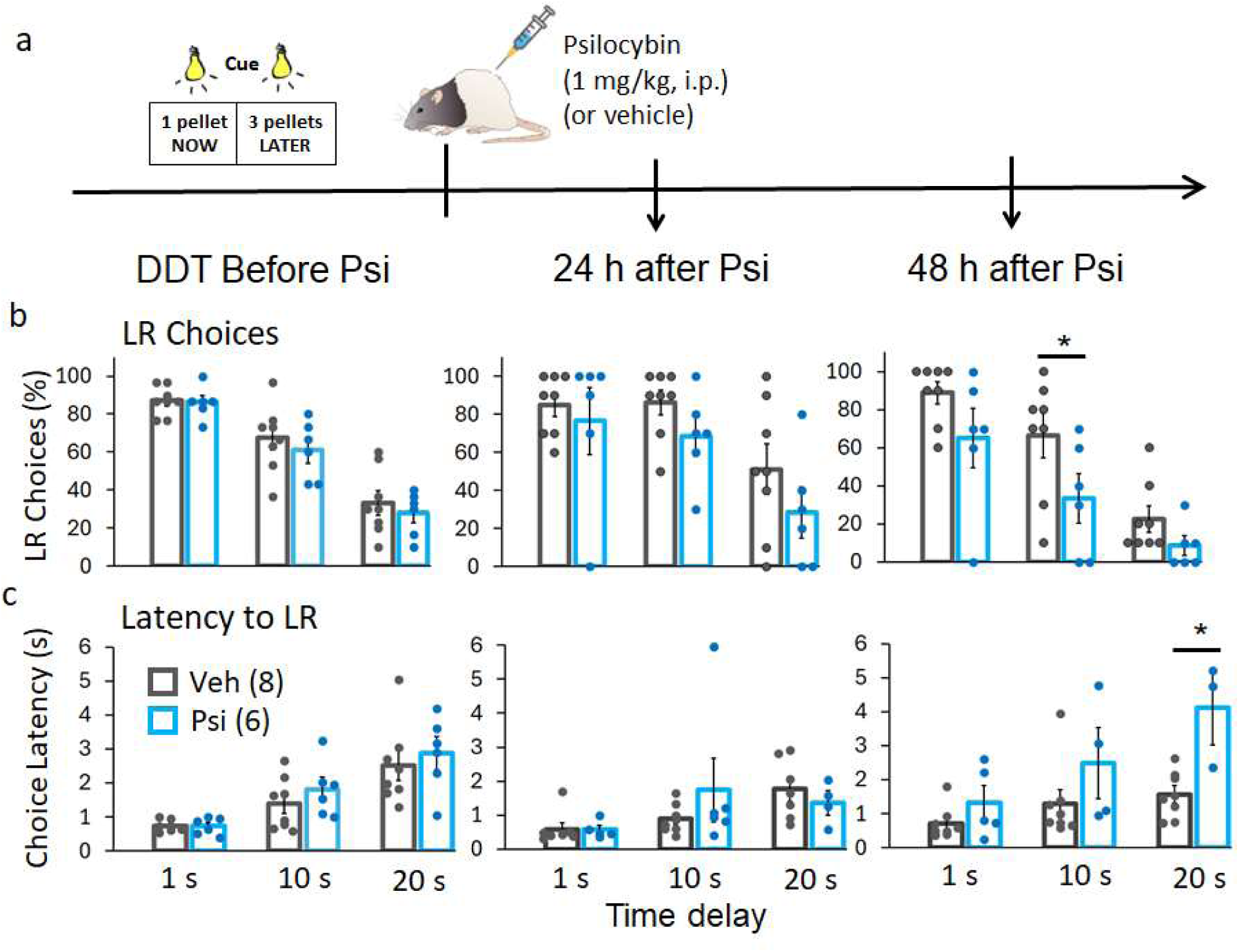
Effects of psilocybin on LR choices and choice latency during Free Choice trials compared to vehicle. **(a)** Timeline for the DDT before and after psilocybin injections. **(b)** LR choices at different delays for LR (1 s, 10 s and 20 s) before psilocybin and 24 and 48 hours after psilocybin compared to vehicle. **(c)** Choice latency at different delays for LR (1 s, 10 s and 20 s) before psilocybin and 24 and 48 hours after psilocybin compared to vehicle. LR choices (%) show a strong trend to decrease after psilocybin compared to vehicle (group: F_(1,12)_= 4.58, p= 0.054, η_p_^2^= 0.28). This decrease was significant at 10 s delay (p= 0.026, η_p_^2^= 0.35). Choice latency (s) decreased after psilocybin compared to vehicle (group: F_(1,8)_= 5.94, p= 0.041, η_p_^2^= 0.43). This decreased depended on the session (session x group: F_(1,8)_= 9.59, p= 0.015, η_p_^2^= 0.54) and was significant at 20 s delay (p= 0.006, η_p_^2^= 0.63). Data are the mean ±SEM. Dots represent individual animals. * p< 0.05 compared to Vehicle after planned comparisons.

**Figure 4:**
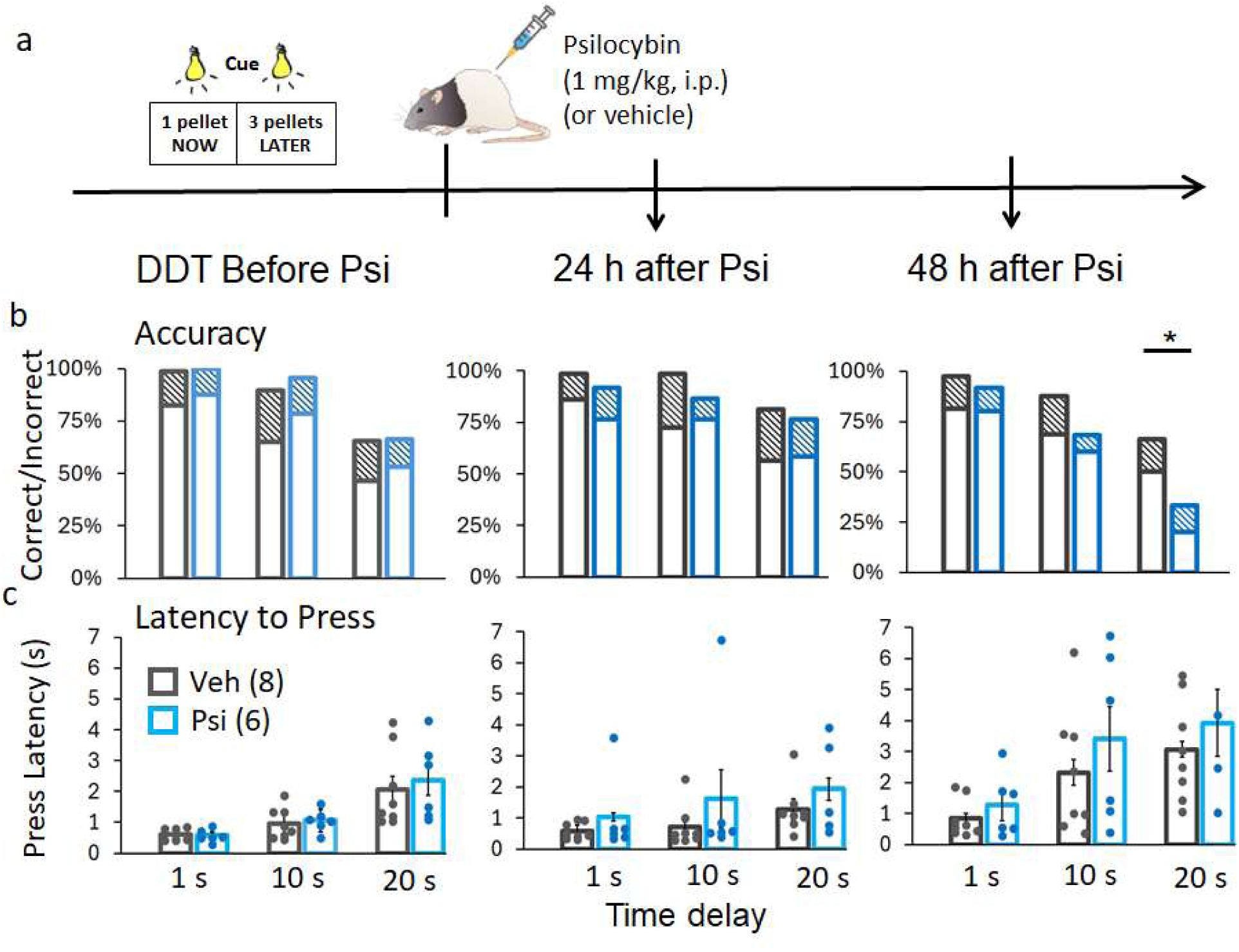
Effects of psilocybin on accuracy and latency to press during Forced Choice trials compared to vehicle. **(a)** Timeline for the DDT before and after psilocybin injections. **(b)** Accuracy (i.e., correct and incorrect presses) at different delays for LR (1 s, 10 s and 20 s) before psilocybin and 24 and 48 hours after psilocybin compared to vehicle. **(c)** Latency to press at different delays for LR (1 s, 10 s and 20 s) before psilocybin and 24 and 48 hours after psilocybin compared to vehicle. Data are the mean ±SEM. Dots represent individual animals. * p< 0.05 compared to Vehicle after planned comparisons.

In Free Choice trials, as shown in **Figure 3b**, psilocybin decreased the number of LR choices compared to vehicle, although this effect did not reach statistical significance (group: F_(1,12)_= 4.58, p= 0.054, η_p_^2^= 0.27). However, planned comparisons demonstrated that this decrease was significant at 10 s delay 48 hours after psilocybin injections (p= 0.026, η_p_^2^= 0.35). The effects of psilocybin on LR choices were not significantly dependent on session (session x group: F_(1,12)_= 0.31, p= 0.584, η_p_^2^= 0.02) or delay (delay x group: F_(2,24)_= 0.40, p= 0.673, η_p_^2^= 0.03). **Figure 3c** shows that psilocybin increased the latency to LR choices compared to vehicle (group: F_(1,8)_= 5.94, p= 0.041, η_p_^2^= 0.43). Planned comparisons showed that this increase was significant at 20 s delay 48 hours after psilocybin injections (p= 0.006, η_p_^2^= 0.63). The effects of psilocybin on choice latency were significantly dependent on session (session x group: F_(1,8)_= 9.59, p= 0.015, η_p_^2^= 0.54) but not delay (delay x group: F_(2,16)_= 2.23, p= 0.140, η_p_^2^= 0.21).

In Forced Choice trials, as shown in **Figure 4b**, psilocybin did not alter accuracy compared to vehicle (group: F_(1,12)_= 1.17, p= 0.30, η_p_^2^= 0.09) even when considering session (session x group: F_(1,12)_= 1.00, p= 0.33, η_p_^2^= 0.07) or delay (delay x group: F_(2,24)_= 0.74, p= 0.485, η_p_^2^= 0.06). However, psilocybin significantly decreased accuracy when considering both session and delay (session x delay x group: F_(2,24)_= 3.42, p= 0.049, η_p_^2^= 0.22). Specifically, planned comparisons demonstrated a significant decrease in accuracy at 20 s delay 48 hours after psilocybin injections (p= 0.026, η_p_^2^= 0.35). **Figure 4c** shows that psilocybin did not alter the latency to press compared to vehicle (group: F_(1,9)_= 0.176, p= 0.68, η_p_^2^= 0.02) even when considering session (session x group: F = 4.40, p= 0.065, η_p_^2^= 0.32) or delay (delay x group: F = 0.98, p= 0.395, η_p_^2^= 0.01). The latency to press in Forced Choice trials increased with higher delays before psilocybin (baseline) (delay: F_(2,18)_= 8.97, p= 0.001, η_p_^2^= 0.52).

Supplemental **Figure 1a** shows that psilocybin did not change the number of SR choices compared to vehicle (group: F_(1,12)_= 0.01, p= 0.901, η_p_^2^= 0.00). The number of SR choices after psilocybin injections was not dependent on session (session x group: F_(1,12)_= 0.46, p= 0.834, η_p_^2^= 0.00) or delay (delay x group: F = 0.48, p= 0.623, η_p_^2^= 0.04). Supplemental **Figure 1b** shows that the number of omissions (i.e., not responding to any lever) during Free Choice trials was not significantly altered by group (group: F_(1,12)_= 2.93, p= 0.112, η_p_^2^= 0.19). Similarly, the number of omissions after psilocybin injections was not dependent on session (session x group: F_(1,12)_= 0.36, p= 0.560, η_p_^2^= 0.03) or delay (delay x group: F_(2,24)_= 0.041, p= 0.960, η_p_^2^= 0.00).

### Effects of psilocybin on PNNs and PVs in the dmPFC and vmPFC

The numerical density of PNNs, cFos and PV neurons immunolabeling was evaluated in both the dmPFC (prelimbic area) and the vmPFC (infralimbic area) of vehicle and psilocybin animals following the behavior experiments. **Figure 5** shows microphotographs indicating superficial and deep medial prefrontal cortex layers and representative labeling of PNNs, cFos and PV neurons. As shown, psilocybin increased numerical densities of triple labelled neurons (PNN+cFos+PV) in the deep layers of both the dmPFC (F-ratio = 6.30, p= 0.03) and the vmPFC (F-ratio = 5.68, p= 0.04). It also shows that psilocybin increased the numerical density of double-labelled PV+cFos neurons in the deep layers of the dmPFC (F-ratio = 5.13, p= 0.04) and decreased the density of single labelled PV neurons in the superficial layers of the dmPFC (F-ratio = 5.95, p= 0.03). Psilocybin also increased the density of PNN+cFos double-labelled neurons in the deep layers of the dmPFC (F-ratio = 5.79, p= 0.03) (Supplemental **Figure 2**). No differences were observed for any of the other marker combinations or for numerical densities of overall PNNs or PV neurons (see Supplemental **Figures 2** and **3**).

**Figure 5:**
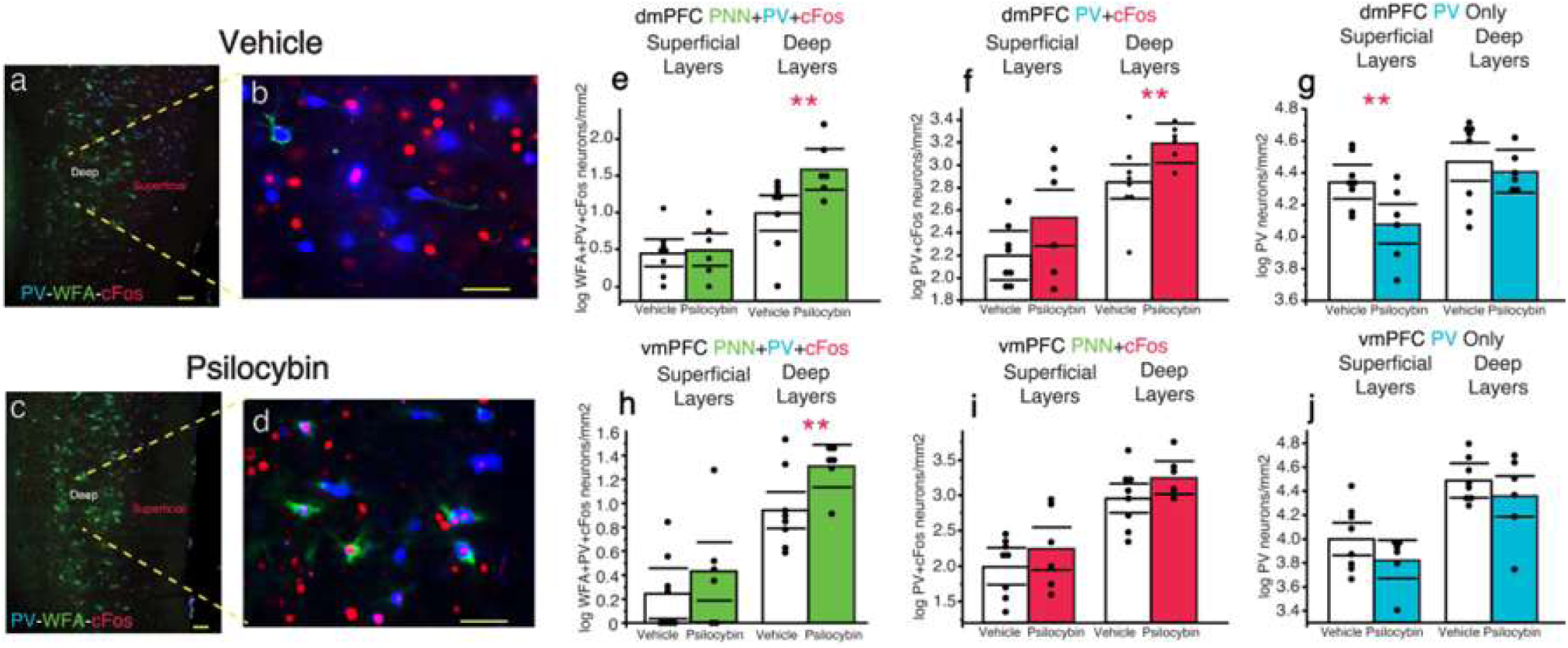
Density of perineuronal nets (PNNs) and parvalbumin (PV) cells positive for cFos in the dmPFC and the vmPFC 48 hours after psilocybin injections. **(a-d)** Microphotographs that highlight triple immunolabeling (PNNs+PV+cFos) in deep cortical layers of the medial prefrontal cortex. Density of PNN+PV+cFos **(e, h)**, PV+cFos **(f, i)** and PV **(g, j)** labelling in both superficial and deep layers of the dmPFC and the vmPFC comparing Vehicle (n = 8) and Psilocybin (n = 6) groups. Bars are the means and dots represent individual animals. Horizontal lines indicate the 95% confidence interval of the samples. **p < 0.01 compared to Vehicle after independent *t* test.

We performed a linear regression analysis to assess whether the density (labelling/mm^2^) of PNNs and PV neurons activity (PNN+cFos+PV labeling) was significantly correlated to the number of LR choices 48 hours after injections, considering both vehicle and psilocybin groups (**Figure 6a**). Changes in LR choices per animal were computed as the average of all delays (1, 10 and 20 s). Pearson’s correlation coefficient showed a significant and inverse correlation between the density of activated interneurons (PV+PNN neurons with cFos) and LR choices in the dmPFC (r= 0.61, p= 0.025), but not in the vmPFC (r= 0.32, p> 0.05). These results aligned with our proposed hypothesis that increased in activity of PV neurons with PNNs contribute to the effects of psilocybin on DDT performance 48 hours after administration (**Figure 6b**).

**Figure 6:**
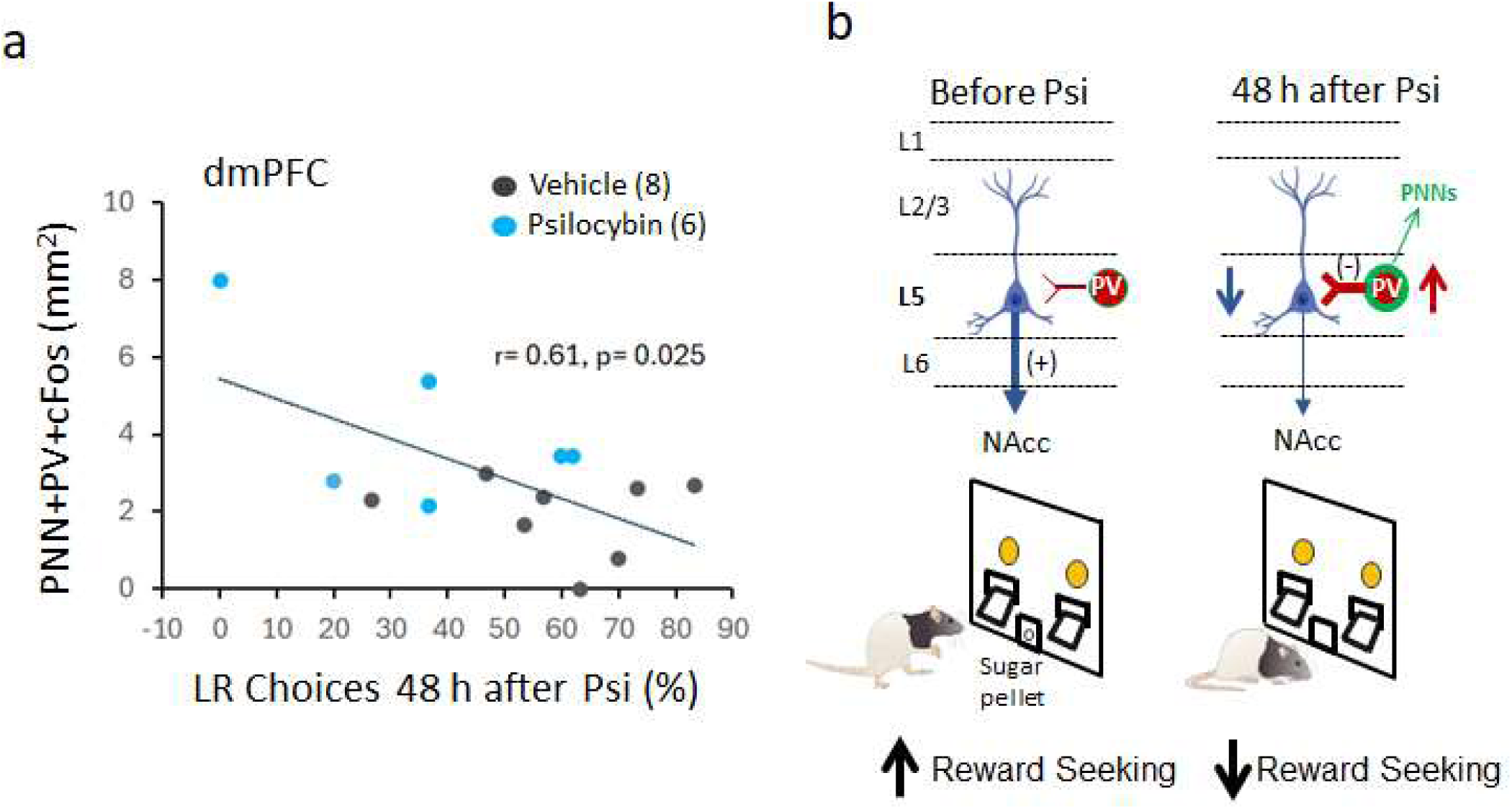
LR choices are negatively correlated to triple (PNN+PV+cFos) labelling in the dmPFC 48 hours after psilocybin. (a) Graph shows the regression analysis including pearson’s correlation coefficient considering both Vehicle and Psilocybin groups. Every dot represents an individual rat. (b) Diagram illustrates the working hypothesis proposed in the current study to integrate both the behavior and immunohistochemistry data shown. PV inhibitory interneurons regulate the activity of pyramidal cells in deep layers of the dmPFC (L5) that project to nucleus accumbens (NAcc) and promote cue-guided reward-seeking behavior (left). Psilocybin increases the activation of PV cells surrounded by PNNs 48 hours after administration. Consequently, we hypothesize that a stronger inhibitory drive mediated by PV interneurons downregulates the activity of dmPFC – NAcc projections which in turns decreases reward-seeking drive during the task (right).

## DISCUSSION

We utilized a value-based reward-seeking task (i.e., DDT) to test the hypothesis that a single dose of psilocybin decreases choice impulsivity in association with changes in activity of PV interneurons surrounded by PNNs in the dmPFC. We found that psilocybin decreased LR lever presses and increased the latency to press during the task 48 hours, but not 24 hours, after administration. Notably, the observed lack of effects at 24 hours is in line with a previous report (Roberts et al. 2023). Critically, the effects at 48 hours were not dependent on delay, and therefore, were not consistent with changes in choice impulsivity. Moreover, we also found increased task-related activation of PV interneurons with PNNs in the deep layers of the dmPFC 48 hours after psilocybin. These results suggest that a single dose of psilocybin decreases the incentive motivation driven by reward cues 48 hours after administration and that these behavioral effects are, in part, mediated by an increased activation of PV inhibitory interneurons in the dmPFC. Furthermore, these results provide novel insights into the long-term mechanisms of action of psilocybin and may help understand its potential role as a treatment for SUDs and other psychiatric disorders.

After DDT training, before psilocybin or vehicle injections, all rats decreased their preference for LR choices and increased their preference for SR choices with higher LR delays during Free Choice trials. These results agree with previous findings, including our own studies (Martinez et al. 2024), using similar DDTs in rats (Saddoris et al. 2015; Sackett et al. 2019; Hernandez et al. 2022) and humans (Kim et al. 2008), and demonstrate that individuals devalue rewards that come with a delay. A decreased preference for LR choices with higher delays is used as an index of increased choice impulsivity and there is an ongoing debate as to how this relates to different aspects of value-based decision making such as reward or time processing (Frost and McNaughton 2017; Levitt et al. 2020; Smith et al. 2023). As expected, we also found an increase in the latency to LR choices with higher delays indicating that a deliberative process is engaged when animals must decide if they will wait more time for preferred rewards. An increased latency to choice also reflects a decreased incentive motivation for the associated reward (Hernandez et al. 2017). Animals also decreased the number of correct responses (Sackett et al. 2019; Martinez et al. 2024) and increased the latency to press with higher delays during Forced Choice trials, which suggests that they also devalued delayed rewards during Forced Choice trials. Notably, they did not change the number of incorrect responses during these trials, which rules out alterations related to attention or task engagement with higher LR delays.

Previous research has shown that a single dose of psilocybin is sufficient to produce sustained behavioral effects and plastic changes in several brain regions including the dmPFC (Shao et al. 2021, 2025; Calder and Hasler 2023; Hatzipantelis and Olson 2024; Anderson and Robinson 2025; Hammo et al. 2025). In the current study, we used one single injection of 1 mg/kg of psilocybin to determine changes in delay discounting behavior and dmPFC markers of PNNs and PV interneurons activity 48 hours after administration. This dose of psilocybin has been shown to modulate cognitive flexibility and attenuate stress-induced behavioral alterations in rodents (Shao et al. 2021; Torrado Pacheco et al. 2023; Anderson and Robinson 2025; Hammo et al. 2025). It also increases synaptic plasticity and gene expression in several areas of the rodent brain, which have been associated with the long-term behavioral effects of psilocybin (Meinhardt et al. 2021; Shao et al. 2021, 2025). Psilocybin acts primarily on 5-HT_2A-2C_ cortical receptors to produce these behavioral and plastic effects (Shao et al. 2021, 2025; Cameron et al. 2023; Hsiao et al. 2025), although 5-HT-independent effects have been also reported (Hesselgrave et al. 2021). In this context, we show here that psilocybin increases head-twitch responses immediately after injection, which agrees with previous studies in rats and confirms the acute activation of 5-HT_2A_ receptors by psilocybin (Willins and Meltzer 1997; Shao et al. 2021) (but see shao et al., 2025).

Most of previous behavioral studies focused on the acute or short-term effects of psilocybin (i.e., from 10 min to 24 hours after administration). Here we assessed performance in the DDT 24 and 48 hours after psilocybin injections. This experimental design allowed us to evaluate the temporal profile of psilocyn effects once it is fully metabolized (Jaster et al., 2025) while seeking for potential long-term actions (i.e., ≥ 48 hours). We found that psilocybin decreased LR choices and increased latency to choice 48 hours, but not 24 hours, after administration. Psilocybin did not change SR choices. Moreover, these effects were independent of delay and therefore are not consistent with psilocybin-induced changes in choice impulsivity. In addition, psilocybin did not change the number of correct (except at 20 s delay) or incorrect responses or the latency to press the lever during Force Choice trials, ruling out general motor or attentional deficits 48 hours after psilocybin. Overall, these results are consistent with psilocybin decreasing motivation for rewards preferentially during Free Choice trials and suggest a long-term effect on brain pathways that regulate cue-driven reward-seeking behavior and incentive motivation.

In line with our findings, recent studies in rats suggest that psilocybin decreases motivation to pursuit rewards in different experimental paradigms. Thus, the 5-HT_2A_ agonist DOI (2,5-Dimethoxy-4-iodoamphetamine) has been shown to decrease cocaine and fentanyl self-administration and motivation for these drugs in male rats (Martin et al. 2021; Salinsky et al. 2025). Similarly, psilocybin has been shown to induce the extinction of opioid rewards (i.e., oxycodone) in male mice by using the conditioned place preference test (Jaster et al. 2025). Interestingly, psilocybin has also been shown to increase reward latency and omissions during a five-choice reaction time task suggesting a decrease motivation to pursuit rewards in this task (i.e., sugar pellets) (Popik et al. 2022). Critically, and differentially from the current study, these studies assessed acute and short-term effects of DOI and psilocybin and therefore do not inform whether the reported effects on reward motivation are long lasting.

The prefrontal cortex regulates reward processing and seeking (Stefanik et al. 2012; Moorman and Aston-Jones 2015; Otis et al. 2017; Laubach et al. 2021; Harris et al. 2025) and is one of the main hubs regulating the effects of psychedelics on the brain (Shao et al. 2021, 2025; Cameron et al. 2023; Siegel et al. 2024). As shown here, psilocybin increased the density of task-induced PV+cFos and PNN+PV+cFos labelling in the dmPFC 48 hours after administration, which indicates an increased activation of PV interneurons in this region of the prefrontal cortex. These results suggest that long-term changes in the function of PV interneurons mediate the observed effects of psilocybin on DDT performance (i.e, decreased reward seeking). Prefrontal PV interneurons are rich in 5-HT_2A_ receptors and therefore, psilocybin could act directly on these neurons to modulate their activity (Willins and Meltzer 1997; De Almeida and Mengod 2007; Andrade 2011; Hsiao et al. 2025). Furthermore, since cFos is significantly increased in PV interneurons with PNNs and PNNs are also modulated by 5HT_2A_ receptors, it is possible that psilocybin enhances the activity of PV interneurons through remodeling PNNs surrounding these neurons (Slaker et al. 2015; Valeri et al. 2023). This possibility fits well with the well-established role of PNNs regulating the activity of PV interneurons (Fawcett et al. 2019; Valeri et al. 2023), although recent research suggests that psilocybin can have long-term cellular and behavioral effects in a manner independent of structural plastic changes (Kramer et al. 2025). Alternatively, prefrontal pyramidal cells and mediodorsal thalamic inputs to dmPFC are also rich in 5HT_2A_ receptors (Andrade 2011; Barre et al. 2016), which opens the possibility that psilocybin indirectly modulates PV interneurons activity in the dmPFC.

Notably, changes in PV interneurons activation were only observed in deep, but not superficial, layers of the dmPFC. Deep layers, specifically layer 5, contain pyramidal neurons projecting to subcortical areas such as the nucleus accumbens, amygdala or thalamus (Gabbott et al. 2005; Otis et al. 2017), which play a key role in reward seeking, choice behavior and incentive motivation (Stefanik et al. 2012; Robinson et al. 2013; Otis et al. 2017; Hernandez et al. 2019; Coley et al. 2021; Pastor and Medina 2021; Wenzel et al. 2023). Deep layers of the dmPFC also contain pyramidal and PV ineterneurons with 5-HT_2A_ receptors and PNNs that control the activity of pyramidal efferent projections (De Almeida and Mengod 2007; Schmitz et al. 2025). Thus, experimental manipulations that change PV interneurons activity or PNNs in the dmPFC modulate reward memory and extinction behavior (Sparta et al. 2014; Slaker et al. 2015). In this context, we also show here that increased labelling of PNNs+PV+cFos in the dmPFC is inversely correlated to decreased LR choices 48 hours after psilocybin administration, which fits well with the idea that the increased activity of PV interneurons mediates, at least in part, the decreased LR choices 48 hours after psilocybin. Considering the inhibitory nature of cortical PV interneurons (Kupferschmidt et al. 2022), our findings support the hypothesis that psilocybin results in the downregulation of dmPFC efferent projections to the nucleus accumbens which promote reward-seeking behavior (Stefanik et al. 2012; Otis et al. 2017). In line with this possibility, recent research highlights the potential role of pyramidal neurons projecting to the nucleus accumbens in mediating the effects of psilocybin on opioid reward extinction (Jaster et al. 2025) and opioid use disorders (Cameron et al. 2023). However, other prefrontal efferent projections have been shown to regulate reward-seeking behavior (Otis et al. 2017; Siciliano et al. 2019; Coley et al. 2021) and therefore future studies are needed to determine how dmPFC pathways may mediate psilocybin-induced changes in reward seeking and motivation.

Together with changes in dmPFC, we also observed an increase in PNN+PV+CFos+ triple labelling in the vmPFC after psilocybin. However, differentially from the dmPFC, these effects did not coincide with significant increases in PV+cFos and PNN+cFos and were not significantly correlated to the number of LR choices 48 hours after psilocybin. Like dmPFC neurons, vmPFC neurons are modulated by psilocybin (Purple et al. 2025; Schmitz et al. 2025) but can have different anatomical connections (Gabbott et al. 2005; Pastor and Medina 2021). Also, behavioral studies suggest that the vmPFC promotes the extinction of reward-seeking behavior (Peters et al. 2009; Warren et al. 2016; Cameron et al. 2019; Ciaramelli et al. 2021; Nett and LaLumiere 2021), while the dmPFC is more directly involved in driving reward seeking (Stefanik et al. 2012; Otis et al. 2017). It should be mentioned that these dmPFC-vmPFC functional differences may depend on the experimental paradigm utilized, and therefore, are still a matter of debate (Moorman and Aston-Jones 2015; Moorman et al. 2015; Gutman et al. 2016; Riaz et al. 2019; Nett and LaLumiere 2021). Overall, our results indicate that psilocybin produces stronger effects on PV interneurons in the dmPFC compared to the vmPFC, which is consistent with a decreased drive to seek out rewards during the DDT.

This study has limitations. First, we only utilized male rats. According to previous studies (Shadani et al. 2024; Jaster et al. 2025), psilocybin and other psychedelic drugs might have different effects depending on sex and therefore it is possible that sex modulates the effects of psilocybin on value-based reward-seeking behavior. Second, the DDT is primarily designed to assess choice behavior more than motivation to pursuit rewards. Nonetheless, the motivation driven by reward cues (i.e., cue light, lever extension) is a key element that guides performance in the DDT. Thus, both a decreased number of LR presses and an increased latency to press are good indexes of decreased motivation for the associated cue. Future studies, however, are needed to identify which features of reward cues (i.e., saliency, value) may be modulated by psilocybin. Finaly, both a relatively small sample size and the evaluation of one or two time points (i.e., 24 and 48 hours), although relevant, are limited to inform about potential effects of psilocybin on reward-related behavior lasting from several days to weeks.

In summary, our study shows that a single dose of psilocybin alters delay-discounting performance in a manner that is consistent with a decreased incentive motivation driven by reward cues. These behavioral alterations coincide with an increased activation of PV interneurons surrounded by PNNs in the dmPFC. Importantly, these behavioral and cellular effects occurred 48 hours after the administration of this psychedelic compound. These findings suggest that psilocybin fine tunes prefrontal inhibitory activity, potentially by remodeling PNNs, to modulate cue-driven reward-seeking behavior. In addition, different from previous studies focusing on acute or short-term effects, our results provide novel insights into the long-term mechanisms by which psilocybin could be useful as a treatment to decrease the escalation of drug intake and relapse in SUDs.

## ACKNOWLEDGEMENTS

The work reported in this article has been supported by the Department of Health and Exercise Science (AD) and the Sally McDonnell Barksdale Honors College (JH and AW), University of Mississippi.

## AUTHOR CONTRIBUTIONS

JH, AW, OA, HP, AD collected the data. HP, BG, AD analyzed the data and wrote the manuscript. AD, HP and BG designed the study. All authors reviewed the manuscript.

## DATA AVAILABILITY

The data and original contributions presented in this study are included in the manuscript and are available upon request to the corresponding author.

## ADDITIONAL INFORMATION

JH and AW contributed equally to this study.

### Competing interest

The authors declare no competing interest.

### ARRIVE

This study is reported in accordance with ARRIVE guidelines.

## Notes

### Competing Interest Statement

The authors have declared no competing interest.

