## Supplemental Figures for "Psilocybin decreases reward-seeking behavior accompanied by increased activity of parvalbumin neurons with perineuronal nets in the medial prefrontal cortex"

**Supplemental Materials**

**Suppl Figure 1:** Effects of psilocybin on SR choices and omissions during Free Choice trials compared to vehicle. **(a)** SR choices at different delays for LR (1 s, 10 s and 20 s) before psilocybin and 24 and 48 hours after psilocybin compared to vehicle. **(b)** Latency to press at different delays for LR (1 s, 10 s and 20 s) before psilocybin and 24 and 48 hours after psilocybin compared to vehicle. Data are the mean ±SEM. Dots represent individual animals.

**Suppl Figure 2:** Density of perineuronal nets (PNNs) only and in cells positive for cFos in the dmPFC and the vmPFC 48 hours after psilocybin injections. Density of PNNs **(a, c)** and PNN+cFos **(b, d)** labelling in both superficial and deep layers of the dmPFC and the vmPFC comparing Vehicle (n = 8) and Psilocybin (n = 6) groups. Bars are the means and dots represent individual animals. Horizontal lines indicate the 95% confidence interval of the samples. $^{**}p < 0.01$ compared to Vehicle after independent *t* test.

**Suppl Figure 3:** Total density of perineuronal nets (PNNs) and PV cells in the dmPFC and the vmPFC 48 hours after psilocybin injections. Density of PNNs **(a)** and PV cells **(b)** labelling in both superficial and deep layers of the dmPFC and the vmPFC comparing Vehicle (n = 8) and Psilocybin (n = 6) groups. Bars are the means and dots represent individual animals. Horizontal lines indicate the 95% confidence interval of the samples.

Suppl Figure 1

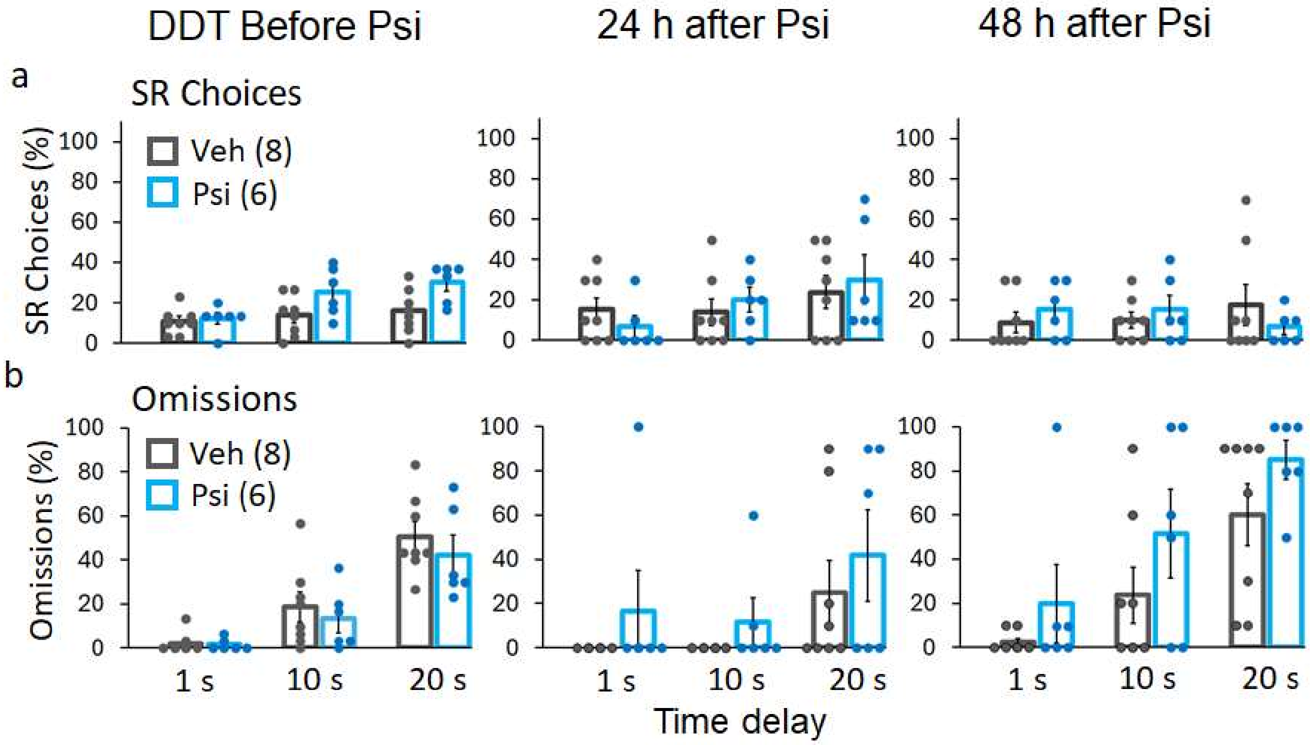

Suppl Figure 2

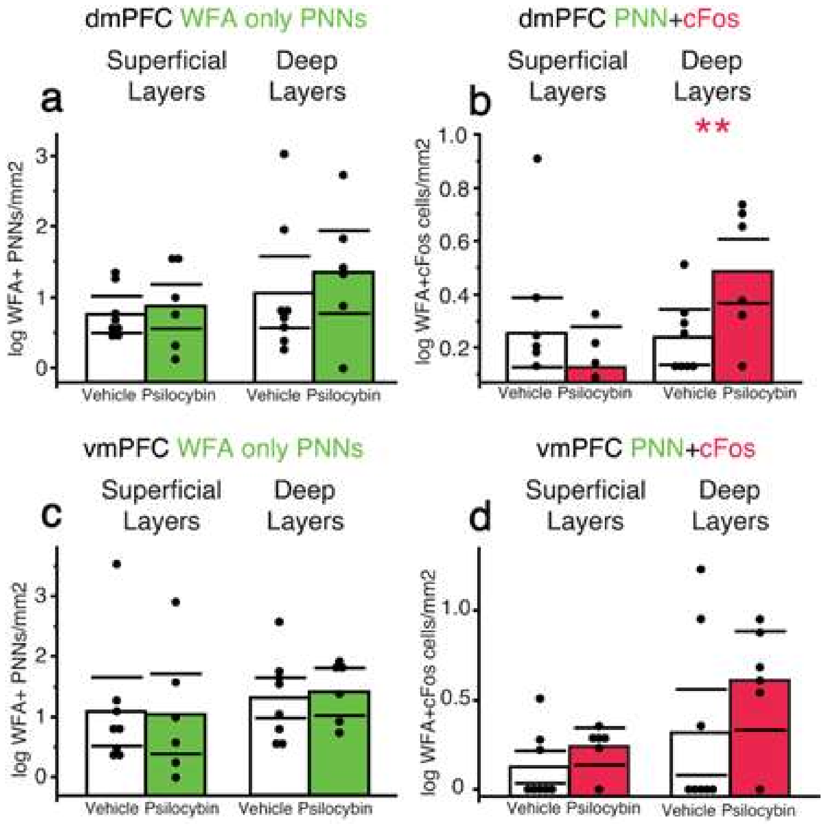

Suppl Figure 3

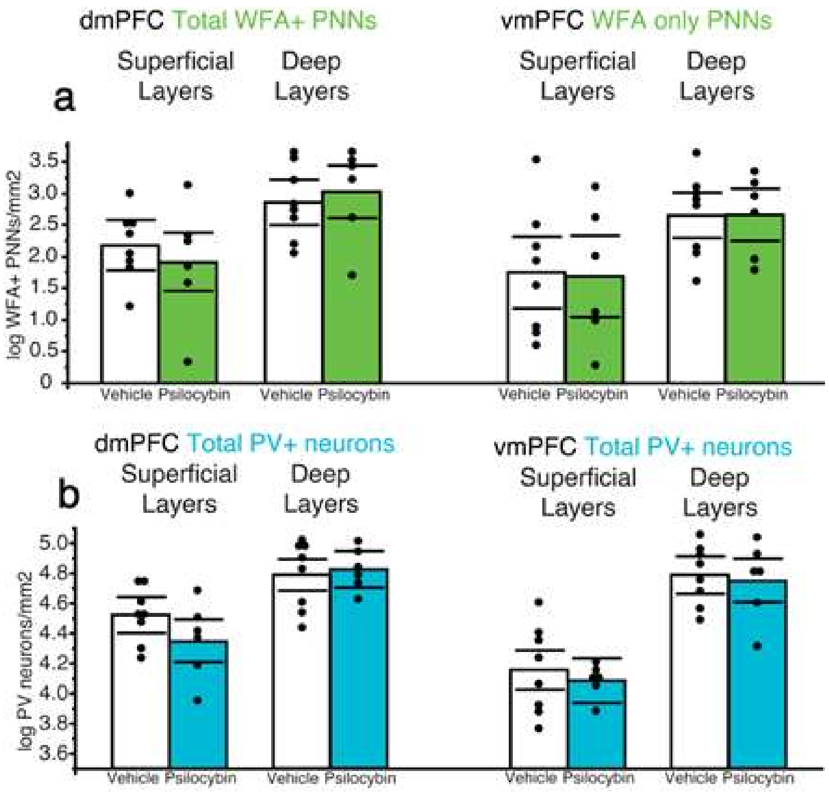
